# Maternal glucocorticoids do not influence HPA axis activity or behavior of juvenile wild North American red squirrels

**DOI:** 10.1101/2020.09.02.280065

**Authors:** Sarah E Westrick, Freya van Kesteren, Stan Boutin, Jeffrey E Lane, Andrew G McAdam, Ben Dantzer

**Author notes:** Corresponding author: Sarah Westrick. Present institution: Department of Evolution, Ecology, and Behavior, University of Illinois at Urbana-Champaign, Urbana, IL, USA.

## Abstract

Environmental factors experienced during development can affect the physiology and behavior of offspring. Maternal glucocorticoids (GCs) may convert environmental cues experienced by the mother into a cue triggering adaptive developmental plasticity in offspring. In North American red squirrels (*Tamiasciurus hudsonicus*), females exhibit increases in GCs when conspecific density is elevated, and selection favors more aggressive and perhaps more active mothers under high density conditions. We experimentally elevated maternal GCs during gestation or early lactation to test the hypothesis that elevated maternal GCs cause shifts in offspring aggression and activity that may prepare them for high density conditions. When offspring were weaned, we measured two behavioral traits (activity and aggression) using a standardized behavioral assay. Because maternal GCs may influence offspring hypothalamic-pituitary-adrenal (HPA) axis activity and HPA axis activity may in turn affect offspring behavior, we also measured the impact of our treatments on offspring HPA axis activity (adrenal reactivity and negative feedback) and the association between offspring HPA axis activity and behavior. Increased maternal GCs during lactation, but not gestation, only slightly elevated activity levels in offspring. Offspring aggression, adrenal reactivity, and negative feedback did not differ between GC-treated and control groups. Offspring with higher adrenal reactivity did exhibit lower aggression, but the relationship between adrenal reactivity and aggression was not affected by treatment with maternal GCs. These results suggest maternal GCs during gestation or early lactation alone may not be a sufficient cue to produce changes in behavioral and physiological stress responses in offspring in natural populations.

**Summary Statement:** We found maternal glucocorticoid levels do not influence offspring personality or HPA axis dynamics in North American red squirrels. Regardless of maternal glucocorticoid treatment, more aggressive squirrels exhibited lower adrenal reactivity.

## Introduction

Maternal effects, or the influence of a mother’s phenotype on those of her offspring, contribute to among-individual phenotypic variation and may induce adaptive developmental plasticity (Lancaster et al., 2007; Reddon, 2012; Rossiter, 1991; Stamps and Groothuis, 2010). For example, gravid female fall field crickets (*Gryllus pennsylvanicus*) exposed to a non-lethal wolf spider (*Hogna helluo*) prior to laying eggs produced more cautious offspring that were more likely to survive in the presence of a lethal wolf spider than control offspring (Storm and Lima, 2010). Maternal glucocorticoids (GCs) have been proposed as one proximate mechanism by which maternal effects shape offspring phenotypes (Kapoor et al., 2008; Meaney, 2001), particularly in regard to the development of their physiological stress response and behavior. Glucocorticoids are responsive to environmental changes, and therefore may act as an indicator that offspring are cued into and respond to with adaptive changes in phenotypes (Del Giudice et al., 2011; Sih, 2011). Similarly, an increase in maternal corticosterone in pregnant female viviparous lizards (*Zootoca vivipara*) improved odds of survival for male offspring (Meylan and Clobert, 2005).

A consistent finding across taxa is that variation in maternal GCs can induce shifts in offspring behavior (Weinstock, 2008). One way in which maternal GCs could lead to adaptive shifts in offspring behavior is through changes in their hypothalamic-pituitary-adrenal (HPA) axis. The HPA axis is a negative feedback system that regulates systemic effector hormones (GCs) (Sapolsky et al., 1985; Spencer and Deak, 2017). Briefly, neural inputs trigger the paraventricular nucleus in the hypothalamus to release corticotropin-releasing factor (CRF) which acts upon the anterior pituitary to release adrenocorticotropic hormone (ACTH) which travels systemically to activate the adrenal cortex to release GCs (cortisol and/or corticosterone) (Packard et al., 2016; Spencer and Deak, 2017). Glucocorticoids are primarily metabolic steroid hormones, but are often studied due to their release in response to stressful situations and adverse environmental conditions (Charmandari et al., 2005; Sapolsky et al., 2000; Spencer and Deak, 2017; Tsigos and Chrousos, 2002). High levels of systemic GCs induce negative feedback by binding to receptors in the hypothalamus and pituitary to return expression of CRF, ACTH, and GC to basal levels after a response to an acute stressor (Sapolsky et al., 1985; Spencer and Deak, 2017).

There are now many laboratory and field studies demonstrating a strong effect of the early life environment and maternal GCs on offspring behavior and HPA axis physiology (Caldji et al., 2011). In an experiment using maternal adrenalectomies and administration of exogenous GCs, Barbazanges et al. (1996) demonstrated the role of excess maternal GCs in mediating the impairment of negative feedback regulation of the offspring’s HPA axis. As summarized in Weinstock (2008), maternal stress can raise GCs and catecholamines, and reduce neural GC receptors in offspring. This reduction in feedback regulation of the HPA axis can alter emotion, cognition, attention, and learning (Meaney, 2001; Weinstock, 2008). For example, increasing GCs of mothers during lactation improved learning and reduced fearfulness of offspring in rats (Catalani et al., 2000), whereas higher GCs in milk of rhesus macaques (*Macaca mulatta*) produced more ‘nervous’ and less ‘confident’ offspring (Hinde et al., 2015). These laboratory studies reveal the potential for changes in maternal GCs to alter offspring behavior, but studies in natural populations are required to understand if the changes in offspring behavior are adaptive.

North American red squirrels (*Tamiasciurus hudsonicus*, hereafter ‘red squirrels’) in the Yukon Territory, Canada, experience among-year fluctuations in the availability of their major food source, seeds from white spruce (*Picea glauca*) trees (Fletcher et al., 2010, 2013; Ren et al., 2017). Red squirrels defend individual territories year-round that each contain a hoard of white spruce cones (Dantzer et al., 2012; Siracusa et al., 2017). Juvenile red squirrels usually must acquire a territory after weaning (usually in the late spring or summer) to survive their first winter (Larsen and Boutin, 1994). The among-year variation in food abundance causes changes in population density such that juvenile red squirrels experience fluctuations in the degree of competition over vacant territories (Taylor et al., 2014). We have previously shown that red squirrels that grow quickly after birth (Dantzer et al., 2013) and those with mothers who were more aggressive and less active (in standardized behavioral assays: Taylor et al., 2014) were more likely to survive under high density conditions. We have also previously found that mothers have elevated GCs during high density conditions (Dantzer et al., 2013; Guindre-Parker et al., 2019) and that elevated GCs during pregnancy result in faster-growing pups (Dantzer et al., 2013, 2020). This suggests that elevations in maternal GCs during pregnancy may induce adaptive increases in offspring growth, but whether or not elevations in maternal GCs cause adaptive increases in offspring aggressiveness is not clear.

In this study, we investigated the effects of elevated maternal GCs on the HPA axis and behavior of their offspring. We asked if changes in maternal GCs induced developmental plasticity in offspring behavior, physiological stress responsiveness, and the interaction between the two by conducting a three-year GC supplementation experiment in a wild population of red squirrels. We tested the hypothesis that elevated maternal GCs induce adaptive shifts in offspring aggressiveness and activity that prepares them for competitive environments. To do so, we increased maternal GCs, which should increase the transfer of maternal GCs to the offspring across the placenta or through milk (Grey et al., 2013; Kulski and Hartmann, 1981; O’Donnell et al., 2009). Our previous studies indicated that elevations in maternal GCs during pregnancy caused increases in offspring postnatal growth, but elevations in maternal GCs during lactation reduced offspring postnatal growth (Dantzer et al., 2020). We therefore investigated how elevations in maternal GCs during pregnancy or lactation affected offspring behavior and HPA axis responsiveness.

We characterized the behavior of offspring from GC-treated mothers and control mothers using standardized behavioral assays that measured offspring activity and aggression towards conspecifics. If maternal GCs induce adaptive plasticity in offspring behavior, we predicted that offspring produced by mothers treated with GCs would be more aggressive and less active. Given that increased exposure to maternal GCs can modify the activity of the HPA axis in offspring (see above), and this may in turn cause changes in offspring behavior, we also quantified the impact of experimentally elevated maternal GCs on offspring HPA axis activity. To do so, we conducted stress challenges that measure the ability of the HPA axis to terminate the physiological stress response (negative feedback following dexamethasone administration: van Kesteren et al. 2019) and to mount a physiological stress response (rise in plasma cortisol concentrations following administration of ACTH: van Kesteren et al. 2019). Based upon previous work in other species (see above), we predicted that offspring produced by mothers treated with GCs would exhibit decreased negative feedback and increased stress responsiveness. As some studies have linked HPA axis activity with behavioral characteristics including aggression and activity (Koolhaas et al., 1999; Westrick et al., 2019), we also examined if there was an association between HPA axis activity and aggression or activity in offspring.

Finally, our previous work showed that activity and aggression in female red squirrels are both phenotypically and genetically positively correlated in red squirrels (Taylor et al., 2012), but selection may favor females who are divergent from that population trend (Taylor et al., 2014). Previous studies in other taxa suggest that elevations in maternal GCs may cause adaptive shifts in phenotypic covariance by either strengthening the degree of covariation among particular traits (Merrill and Grindstaff, 2015) or lessening the overall degree of phenotypic covariance (Careau et al., 2014). We therefore investigated if our GC treatments affected the phenotypic correlation between activity and aggression. We predicted that juveniles produced by mothers with elevated GCs would exhibit a significantly reduced positive correlation between aggression and activity compared with offspring from control mothers.

## Materials and Methods

### Study population

We studied red squirrels in two different study areas on the traditional lands of the Champagne and Aishihik First Nations in the Yukon Territory, Canada (61**°**N, 138**°**W). North American red squirrels of both sexes are highly territorial and defend their food cache year-round (Dantzer et al., 2012; Siracusa et al., 2017). To identify adults, we used unique alphanumeric stamped ear tags (National Band and Tag Company, Newport, KY, USA) and unique combinations of colored wire threaded through both ears tags. We used live trapping (Tomahawk Live Trap, Tomahawk, WI, USA) to monitor reproductive status by abdominal palpation to detect fetuses and manual milk expression to detect lactation (McAdam et al., 2007). Upon identifying a lactating female, we collared her with a VHF radio transmitter (Holohil PD-2C, 4 g, Holohil Systems Limited, Carp, Ontario, Canada),and used telemetry to locate the nest containing her pups. We briefly removed the pups from the nest soon after birth, and again ∼25 days post-parturition in order to identify sexes, record masses to calculate post-natal growth, and ear-tag pups with alphanumeric tags and unique combinations of colored disks for identification after emergence at ∼35 days. Red squirrel pups are completely weaned around 70 days (Boutin and Larsen, 1993). For more details on the general population monitoring methods see McAdam et al. (2007). All work was conducted under animal ethics approvals from the University of Michigan (PRO00005866).

### Glucocorticoid supplementation experiment

To stimulate chronic increases in GCs, we treated pregnant or lactating females with exogenous GCs mixed with all natural peanut butter using methods described in Dantzer et al. (2013) and van Kesteren et al. (2019). Control females received the same peanut butter treatments without GCs. To treat individuals with GCs daily, we provisioned them with small amounts of peanut butter (∼8 g) and wheat germ (∼2 g) mixed with dissolved hydrocortisone [H4001, Sigma Aldrich]. We first dissolved the hydrocortisone in 1 mL of 100% ethanol before mixing with 5 mL of peanut oil. We let the emulsion sit overnight to evaporate the ethanol. We combined the peanut oil with 800 g peanut butter and 200 g wheat germ, weighed out individual doses (∼10 g), placed each dose in individual containers, and stored doses at -20 °C until needed for provisioning the squirrels. Control treatments were made in exactly the same manner but did not include hydrocortisone in the peanut oil.

On the territory of each squirrel in our experiment, we hung a 10.5 L bucket with two holes cut into its sides. Each bucket was covered with a lid and hung ∼7-10 m off the ground in the center of the squirrel’s territory. We placed individual peanut butter treatments in the buckets for provisioning each day. We randomly assigned squirrels to either the control treatment (8 g all-natural peanut butter, 2 g wheat germ, no GCs) or GC treatment (8 g all-natural peanut butter, 2 g wheat germ, 8 or 12 mg of GCs). We fed all pregnancy treated squirrels 8 mg of GCs/day and lactation treated squirrels either 8 or 12 mg of GCs/day. We selected these dosages of GCs to keep GCs within physiological levels, based on previous studies in red squirrels (Dantzer et al., 2013; van Kesteren et al., 2019) and laboratory rats (Casolini et al., 1997; Catalani et al., 2002). Wilcoxon Rank Sum tests on the four response variables (two HPA axis measurements and two behavioral traits) showed no significant differences between 8 and 12 mg of GCs/day in the lactation treatment group (Table S1). Therefore, we combined the 8 and 12 mg treatments into one GC treatment group to increase statistical power (Dantzer et al., 2020).

To examine whether the timing of an increase in maternal GCs produced unique changes in offspring phenotypes, we treated breeding female squirrels *either* during late pregnancy *or* during early lactation. In the pregnancy treatment groups, we aimed to treat mothers for 20 days starting approximately 15 days prior to birth (20 days after conception) until 5 days after birth. Due to variation in detecting the precise stage of pregnancy via palpation, we actually treated mothers from 10.8 ± 0.7 (mean ± SD) days prior to birth to 4.6 ± 0.4 days after birth (actual treatment length: 16.3 ± 0.6 days). In the lactation treatment groups, we aimed to treat mothers for 10 days starting 5 days after birth until 15 days after birth. We actually treated mothers during lactation 5.1 ± 0.2 days post-parturition to 14 ± 0.3 days post-parturition (actual treatment length: 10 ± 0.1 days). This experimental design resulted in four treatment groups which we will refer to as: preg control, preg GC, lac control, and lac GC (see Table 1 for sample sizes). For more detailed information about this manipulation and how it impacts circulating levels of cortisol (the major glucocorticoid in red squirrels) in plasma, see van Kesteren et al. (2019).

**Table 1.** Sample Sizes. Number of red squirrel mothers supplemented, litters produced, and pups tested within each treatment group. Two mothers are represented in alternating treatment groups (GC and control) during pregnancy in different years. Not all juvenile red squirrels underwent an OF/MIS trial, so the sample sizes for analyses of HPA axis dynamics without any behavioral variables (A) included a few more squirrels than the analyses of HPA axis dynamics and behavior (‘activity’ and ‘aggression’) (B).

**A) HPA axis dynamics by treatment group**
|  | **Pregnancy Treatments** | **Lactation Treatments** |
| --- | --- | --- |
| GC peanut butter<br>(8 or 12 mg hydrocortisone) | 13 mothers supplemented<br>13 litters produced<br>20 pups tested | 8 mothers supplemented<br>8 litters produced<br>13 pups tested |
| control peanut butter<br>(0 mg hydrocortisone) | 13 mothers supplemented<br>13 litters produced<br>21 pups tested | 9 mothers supplemented<br>9 litters produced<br>13 pups tested |

**B) HPA axis dynamics and behavior by treatment group**
|  | **Pregnancy Treatments** | **Lactation Treatments** |
| --- | --- | --- |
| GC peanut butter<br>(8 or 12 mg hydrocortisone) | 12 mothers supplemented<br>12 litters produced<br>16 pups tested | 7 mothers supplemented<br>7 litters produced<br>12 pups tested |
| control peanut butter<br>(0 mg hydrocortisone) | 12 mothers supplemented<br>12 litters produced<br>18 pups tested | 9 mothers supplemented<br>9 litters produced<br>11 pups tested |

### Behavioral trials

We live-trapped offspring from our experimental females around the age of weaning (mean ± SD = 67.79 days old ± 3.76; weaning age is ∼70 days old: Boutin and Larsen 1993). Using a canvas handling bag, we weighed the juvenile squirrels before transferring them to our behavioral assay arena. For our open-field and mirror image stimulation trials, we used the same white corrugated plastic arena (60 × 80 × 50 cm) with a clear acrylic lid as described in previous studies in this study system (Boon et al., 2007; Boon et al., 2008; Kelley et al., 2015; Taylor et al., 2012, 2014). We recorded the squirrel’s behavior using a digital video camera for later scoring. For the open-field trial, squirrels were in the open arena for 7 min. This also served as the acclimation period for the following mirror image stimulation trial which lasted 5 min after the mirror (45 × 30 cm) on one side of the arena was exposed.

We used JWatcher (Blumstein and Daniel, 2007) to manually score the videos using the same ethogram used in previously published studies of red squirrel personality (Boon et al. 2007, 2008; Taylor et al. 2012, 2014; Westrick et al. 2019). Observers (n = 4) were blind to the treatment group of the individual. In our analyses, we only included behaviors that previously showed high inter-observer reliability (Taylor et al., 2012).

### HPA axis hormone challenges

We performed HPA axis hormone challenges by administering dexamethasone (DEX; a GC receptor antagonist) and ACTH as previously described to offspring from our experimental females (van Kesteren et al., 2019). Briefly, DEX binds to the GC receptors to induce negative feedback of the HPA axis, primarily through acting on the anterior pituitary (De Kloet et al., 1975), which downregulates circulating GC levels, while ACTH acts upon the adrenals to upregulate the production of GCs. We began by collecting a blood sample from a rear toenail using heparinized microcapillary tubes (described in van Kesteren et al. 2019; Dantzer et al. 2020) for measurement of initial concentrations of total cortisol circulating in plasma. It is important to note that this sample was taken after trapping, handling, recording through a behavioral trial, and transporting individuals from their natal or recently claimed territories to our field station. Due to this substantial amount of handling and disturbance, we consider the initial sample levels of cortisol as stress-induced, though the length of time between trapping and the first bleed was variable. We then injected 3.2 mg/kg of dexamethasone intramuscularly in the squirrel’s upper rear leg. We released the squirrel back into the live-trap and waited 1 hr before taking another blood sample (DEX bleed). Next, we injected 4 IU/kg of ACTH intramuscularly in the alternate upper rear leg. We kept the squirrel in the live-trap before taking blood samples 30 mins (ACTH 30) and 1 hr post-injection (ACTH 60). We have previously shown that these dosages of dexamethasone and ACTH are sufficient to decrease or increase circulating plasma cortisol concentrations in adult red squirrels (van Kesteren et al., 2019). We kept all blood samples on wet ice during the challenge. At the conclusion of the challenge, we separated the plasma via centrifugation and then froze the samples at -80 °C.

To quantify total plasma cortisol concentrations, we used an ImmuChem coated tube cortisol radioimmunoassay (RIA) kit (MP Biomedicals) following the manufacturer’s instructions with minor modification of sample and tracer volumes, and ran samples in duplicate, when possible. To run as many duplicates as possible with our small plasma volumes, we used 12.5 µl of sample and 500 µl of tracer. We ran 87% of samples in duplicate. On rare occasions, we were unable to collect enough blood to quantify total plasma cortisol at every time point (initial sample n = 5, DEX bleed n = 3, ACTH 30 min n = 6, ACTH 60 min n = 4). To maximize the number of squirrels included in our study, we used the global mean value of total plasma cortisol for that respective time point for these missing time points. Due to the large number of samples, we ran RIAs on four different days. Across all four assays, our average standard and sample intra-assay CVs were 9.5%, our average intra-assay CVs for red squirrel plasma samples was 9.3%, and our average inter-assay CVs for the five standards provided (10, 30, 100, 300 and 1000 ng/ml cortisol) was 14%. The experimenters conducting the HPA axis challenges were not blind to maternal treatments due to the same researchers provisioning the mothers and trapping the squirrels but were blind to the results of the behavioral trials. The experimenters conducting the RIAs were blind to both the maternal treatments and results of the behavioral trials.

### Statistical analyses

We ran all statistical analyses in R version 3.5.2 (R Core Team, 2016). Using the R package ‘ade4’ version 1.7-10 (Dray and Dufour, 2007), we ran two distinct principal components analyses with correlation matrices (one for open-field behaviors and one for mirror-image stimulation behaviors) to reduce behavioral variables down to one major component for each assay. Based on the loadings (Table 2), we interpreted the first component of the open-field trial as ‘activity’, explaining 30% of variation in open-field behaviors in our data set. We interpreted the first component of the mirror image stimulation trial as ‘aggression’, explaining 50% of variation in mirror image stimulation behaviors in our data set. Previous studies in this system have used the same methods to analyze open-field and mirror image stimulation trials, and also used the same interpretation for the first component for the open-field trial and mirror image stimulation trial (Boon et al., 2007, 2008; Cooper et al., 2017; Kelley et al., 2015; Taylor et al., 2012, 2014; Westrick et al., 2019). All subsequent analyses used the individual scores calculated from the principal component loadings for each trial (Table 2). Higher ‘activity’ scores mean the squirrel spent more time walking, jumping etc. Higher ‘aggression’ scores mean the squirrel attacked the mirror more often and spent more time in front of the mirror than lower scoring squirrels (Table 2).

**Table 2.** PCA loadings for behaviors scored in the open-field trials (OF) and mirror image stimulation trials (MIS) PCA loadings for juvenile red squirrel behaviors with high inter-observer reliability for the first PCA component for both the open-field and mirror image stimulation trials (Taylor et al., 2012). We used PCA loadings over 0.2 for the interpretation of the component. We calculated ‘activity’ and ‘aggression’ scores from these loadings.

| OF Behavior | Loading |
| --- | --- |
| time spent walking | 0.59 |
| time spent hanging | 0.33 |
| chewing or scratching | 0.19 |
| number of jumps | 0.79 |
| hole head dips | 0.29 |
| time spent grooming | -0.47 |
| not moving | -0.81 |
| Proportion of Variance | 0.30 |

| MIS Behavior | Loading |
| --- | --- |
| time spent in front of arena | 0.77 |
| number of attacks | 0.48 |
| time spent in back of arena | -0.68 |
| latency to attack | -0.75 |
| latency to approach | -0.80 |
| Proportion of Variance | 0.50 |

We calculated the adrenal responsivity, or the net integrated response of cortisol over the 60 mins post-ACTH injection, as the area under the curve (AUC) from the DEX bleed to ACTH 60 using the natural cubic spline interpolation (ACTH AUC). Area under the curve is used in other mammalian and avian study systems as a measure of the integrated adrenocortical response to ACTH (Heidinger et al., 2008; Ingram et al., 1997; Janssens et al., 1994; Rich and Romero, 2005; Saltzman et al., 2000). Based on a recent review about calculating HPA negative feedback after a DEX injection (Lattin and Kelly, 2020), we calculated the relative decrease in cortisol from the initial sample bleed to the DEX bleed (DEX response).

We used the R package ‘lme4’ version 1.1-19 (Bates et al., 2015) to fit linear mixed-effects models and estimated *p*-values using the R package ‘lmerTest’ version 3.0-1 (Kuznetsova et al., 2016). We used the R package ‘multcomp’ version 1.4-8 (Hothorn et al., 2008) to run a Tukey post-hoc comparison following a linear mixed-effects model of plasma cortisol levels at each of the four sampling time points. For each response variable (HPA axis and behavior variables), we fit separate models for pregnancy and lactation treatment groups. To control for the variability in the number of doses mothers in the pregnancy treatment group received (mean = 16 ± 3.59 days), we included maternal treatment length in all pregnancy models. Treatment length among mothers in the lactation groups did not vary considerably (mean = 10 ± 0.49 days), therefore we did not include this in the lactation models. We compared the HPA response variables (ACTH AUC and DEX response) between juveniles in GC-treated groups to the appropriate control groups using linear mixed-effects models. In the pregnancy models to predict ACTH AUC, we included treatment group (GC or control), sex, post-DEX injection plasma total cortisol concentration, maternal treatment length (standardized across all data for all analyses), treatment year (categorical), and age of the juvenile (standardized across all data for all analyses) as fixed effects. In the pregnancy models to predict the response to DEX, we included treatment group, maternal treatment length, sex, and age. We removed year of treatment from this model due to singular fit, indicating the model was overfit, when it was included. Since we included multiple pups from the same litter, we included a litter identity (ID) as a random effect. For the lactation model predicting ACTH AUC, we included treatment group, sex, post-DEX injection plasma total cortisol concentration, treatment year, and age as fixed effects and excluded litter ID as a random effect. In the lactation model predicting response to DEX, we included treatment group, sex, treatment year, and age as fixed effects, and litter ID as a random effect.

To assess the impact of maternal GCs on behavior and the relationship between HPA and behavior, we fit separate linear mixed-effects models for activity and aggression. In our pregnancy treatment activity model, we included ACTH AUC, response to DEX, treatment group, sex, maternal treatment length, treatment year, and age as fixed effects. In our lactation treatment activity model, we included ACTH AUC, response to DEX, treatment group, treatment year, and age as fixed effects, but did not include sex due to singular fit when it was included. We included litter ID in both models as a random effect. For our pregnancy treatment aggression model, we fit a linear model predicting aggression including ACTH AUC, response to DEX, treatment group, sex, maternal treatment length, treatment year, and age as fixed effects. Due to overfitting, we were unable to fit a linear mixed-effects model including litter ID as a random effect for this model. The maximum number of pups from each litter in our sample is two (n = 7 litters with two pups out of 16 total litters). For the lactation treatment aggression linear mixed-effects model, we included ACTH AUC, response to DEX, treatment group, sex, treatment year, and age as fixed effects and litter ID as a random effect.

To assess the impact of maternal treatment on the relationship between activity and aggression, we fit linear mixed-effects models with activity as the response variable and fixed effects including the interaction between aggression and maternal treatment group, treatment year, and sex with litter ID as a random effect. For the pregnancy model, we included an additional fixed effect of maternal treatment length. We excluded age as a variable in both models due to singular fit. To detect any collinearity in the predictors included in our models, we used R package ‘car’ version 3.0-2 (Fox and Weisberg 2011) to assess the variance inflation factors. We found GVIF^(1/(2xDF))^ < 2 for all predictors across all models. Due to small sample sizes, we did not include an interaction between sex and treatment in any models.

## Results

On average, all treatment groups responded to DEX and ACTH as expected with, respectively, a decrease in plasma cortisol concentrations, and subsequent increase in plasma cortisol concentrations (Figure 1). A simple linear mixed model with Tukey post-hoc comparisons showed plasma cortisol concentrations did not differ between the initial handling-stress induced sample and the ACTH 30 min bleed (ACTH 30 - initial sample: ß = -4.77, z = -2.03, *p* = 0.18) and were lowest at the DEX bleed, 1 hr after the injection of DEX (DEX bleed - initial sample: ß = -36.78, z = -15.67, *p* < 0.001; ACTH 30 – DEX: ß = -32.01, z = 13.66, *p* < 0.001; ACTH 60 - DEX: ß = 27.08, z = 11.55, *p* < 0.001). Plasma cortisol concentrations did not differ significantly between ACTH 30 min and ACTH 60 min bleeds (ACTH 60 – ACTH 30: ß = -4.93, z = -2.10, *p* = 0.15). One individual (out of 57 total individuals) from the preg control treatment group did not respond to DEX and was excluded from further analyses.

**Figure 1.**
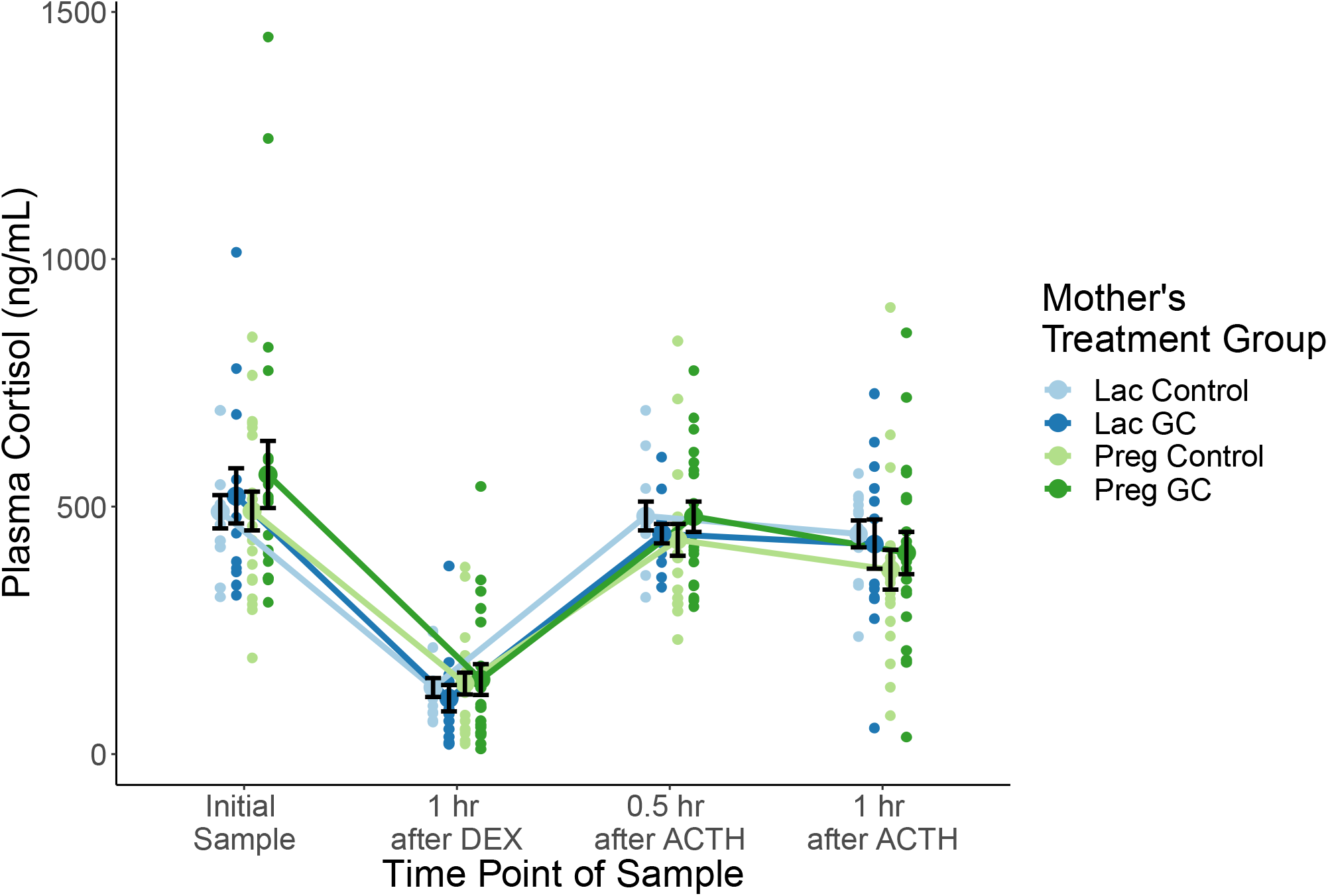
HPA axis hormone challenge curves by treatment group. Plasma cortisol concentrations (ng/mL) of juvenile red squirrels plotted across the DEX/ACTH hormone challenge time series. Blue lines and points indicate juveniles from lactation treatments (lac control n = 12 juvenile squirrels, lac GC n =13 juvenile squirrels) and green lines and points indicate juveniles from pregnancy treatments (preg control n = 20 juvenile squirrels, preg GC n = 20 juvenile squirrels). Data are staggered at each timepoint by mother’s treatment group for ease of visualizing. Black bars indicate the standard error around the mean for each group at each timepoint.

### Effect of maternal glucocorticoids on HPA axis activity

Juveniles from mothers treated with exogenous GCs during pregnancy or lactation did not differ in their adrenal response to ACTH, as measured by AUC, compared to controls (pregnancy treatment: ß = 8.54, *p* = 0.23; lactation treatment: ß = -8.45, *p* = 0.32; Table 3; Figure 2). There were no sex differences in the response to ACTH (Table 3). Older pups in the lactation treatment group (but not in the pregnancy group) had smaller responses to ACTH (ß = -7.72, *p* = 0.015). Likewise, treatment length for pregnancy treatments did not explain variation in ACTH AUC (ß = -1.85, *p* = 0.74; Table 3A). We found no effect of year on ACTH AUC across the three years of this experiment (Table 3).

**Table 3.** Full results of HPA axis dynamics models. We ran four distinct models testing the effect of maternal GCs on the HPA axis dynamics in juvenile red squirrels. We ran separate models for the (a) pregnancy and (b) lactation treatment groups and the two different aspects of the HPA axis dynamics (area under the curve and negative feedback). We standardized treatment length (days) across all data for the pregnancy group and standardized age of pup (days) across all data. The comparison group for categorical variables is control treatment females from 2015. Bold font indicates statistical significance of *p* < 0.05.

**A) Pregnancy treatments**
| Response Variable | Fixed Effect | $\beta$ | SE | t | p-value |
| --- | --- | --- | --- | --- | --- |
| **Area under the curve (DEX to ACTH 60)** |  |  |  |  |  |
| n = 38 pups | **Intercept** | **75.25** | **10.20** | **7.38** | **< 0.0001** |
| n = 25 litters | Treatment |  |  |  |  |
|  | preg GC | 8.54 | 6.82 | 1.25 | 0.23 |
|  | Sex |  |  |  |  |
|  | males | -3.00 | 6.68 | -0.45 | 0.66 |
|  | **DEX cortisol (ug/dL)** | **1.13** | **0.24** | **4.67** | **< 0.0001** |
|  | Treatment length (days) | -2.22 | 5.33 | -0.42 | 0.68 |
|  | Year |  |  |  |  |
|  | 2016 | -23.77 | 12.40 | -1.92 | 0.07 |
|  | 2017 | -22.05 | 12.50 | -1.76 | 0.09 |
|  | Age of pup (days) | -2.07 | 4.48 | -0.46 | 0.65 |
|  | Random effect: | Variance |  |  |  |
|  | Litter ID | 84.3 |  |  |  |
|  | Residual | 270.1 |  |  |  |
| **Relative decrease from initial handling-stressed sample to DEX (%)** |  |  |  |  |  |
| n = 40 pups | **Intercept** | **67.51** | **5.40** | **12.50** | **< 0.0001** |
| n = 26 litters | Treatment |  |  |  |  |
|  | preg GC | 9.77 | 6.48 | 1.51 | 0.14 |
|  | Sex |  |  |  |  |
|  | males | -9.09 | 6.65 | -1.37 | 0.18 |
|  | Treatment length (days) | 6.65 | 3.89 | 1.71 | 0.10 |
|  | Age of pup (days) | -0.57 | 3.77 | -0.15 | 0.88 |
|  | Random effect: | Variance |  |  |  |
|  | Litter ID | 21.45 |  |  |  |
|  | Residual | 360.76 |  |  |  |

**B) Lactation treatments**
| Response Variable | Fixed Effect | $\beta$ | SE | t | p-value |
| --- | --- | --- | --- | --- | --- |
| **Area under the curve** |  |  |  |  |  |
| **(DEX to ACTH 60)** |  |  |  |  |  |
| n = 25 pups | **Intercept** | **63.11** | **9.50** | **6.64** | **< 0.0001** |
| n = 17 litters | Treatment |  |  |  |  |
|  | lac GC | -8.45 | 8.13 | -1.04 | 0.32 |
|  | Sex |  |  |  |  |
|  | males | 4.39 | 5.41 | 0.81 | 0.43 |
|  | **DEX cortisol (ug/dL)** | **1.59** | **0.20** | **7.80** | **0.00035** |
|  | Year |  |  |  |  |
|  | 2016 | -8.04 | 13.55 | -0.59 | 0.56 |
|  | 2017 | 0.93 | 8.63 | 0.11 | 0.92 |
|  | **Age of pup (days)** | **-7.72** | **2.60** | **-2.96** | **0.015** |
|  | *Random effect:* | *Variance* |  |  |  |
|  | Litter ID | 193.91 |  |  |  |
|  | Residual | 63.06 |  |  |  |
| **Relative decrease from initial handling-stressed sample to DEX (%)** |  |  |  |  |  |
| n = 25 pups | **Intercept** | **64.35** | **16.51** | **3.90** | **< 0.001** |
| n = 17 litters | Treatment |  |  |  |  |
|  | lac GC | -0.11 | 9.10 | -0.01 | 0.99 |
|  | Sex |  |  |  |  |
|  | males | -0.49 | 8.20 | -0.06 | 0.95 |
|  | Year |  |  |  |  |
|  | 2016 | -8.37 | 13.43 | -0.62 | 0.55 |
|  | 2017 | -4.41 | 8.53 | -0.52 | 0.62 |
|  | **Age of pup (days)** | **-6.46** | **4.04** | **-1.60** | **0.13** |
|  | *Random effect:* | *Variance* |  |  |  |
|  | Litter ID | 14.45 |  |  |  |
|  | Residual | 330.62 |  |  |  |

**Figure 2.**
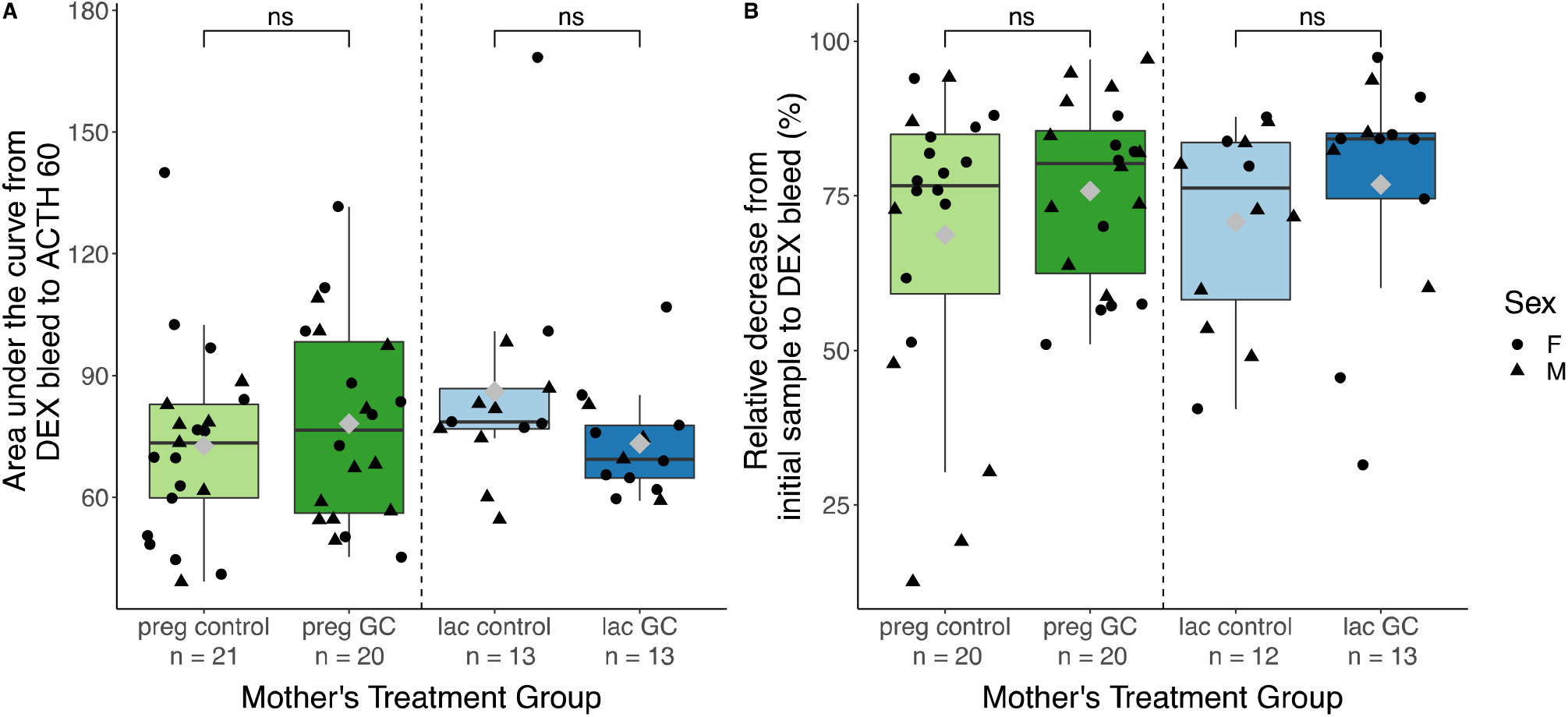
Effect of treatment group on HPA axis dynamics. A) Adrenal reactivity (response to ACTH) and B) negative feedback (response to dexamethasone) of juvenile red squirrels were not impacted by maternal treatment with exogenous GCs (Table 3). Box plots indicate the lower quartile, median, and upper quartile. Whiskers indicate the range of the data. Grey diamonds indicate the group means. Brackets above box plots indicate comparisons between corresponding control and GC treatment groups (ns = not significant, *p* > 0.05). Data points represent individual juvenile red squirrels (preg control n = 21, preg GC n = 20, lac control n = 13, lac GC n = 13). The shape of the data point indicates the sex of the individual (circle = female red squirrel, triangle = male red squirrel).

Juveniles from mothers treated with exogenous GCs during pregnancy or lactation also did not differ in the magnitude of their negative feedback response to DEX compared to controls (pregnancy treatment: ß = 9.77, *p* = 0.14; lactation treatment: ß = -0.11, *p* = 0.99; Table 3; Figure 2). The sex and age of the juvenile did not contribute significantly to variation in negative feedback, and neither did treatment length for pregnancy treatments (Table 3). We found no differences in negative feedback among years of the experiment (Table 3).

### Effect of maternal glucocorticoids on behavioral traits

Among juveniles from both preg GC and preg control mothers, more active individuals exhibited a lower adrenal response to ACTH (ACTH AUC) than less active individuals (ß = - 0.02, *p* = 0.0058; Table 4A; Figure 3). However, juveniles from lac GC and lac control mothers did not exhibit any relationship between ACTH AUC and activity (ß = 0.01, *p* = 0.66; Table 4B; Figure 3). We saw a non-significant trend for juveniles from lac GC mothers to have slightly higher activity levels than control individuals (ß = 1.17, *p* = 0.07; Table 4B; Figure 4). However, preg GC offspring were not significantly different than preg control offspring in terms of activity levels (Table 4A; Figure 4). Again, sex and age of the juvenile did not contribute to variation in activity (Table 4). We found that the negative feedback response to DEX did not predict activity in either pregnancy or lactation treatments (Table 4; Figure 3). We found no differences in activity among years of the experiment (Table 4).

**Table 4.**
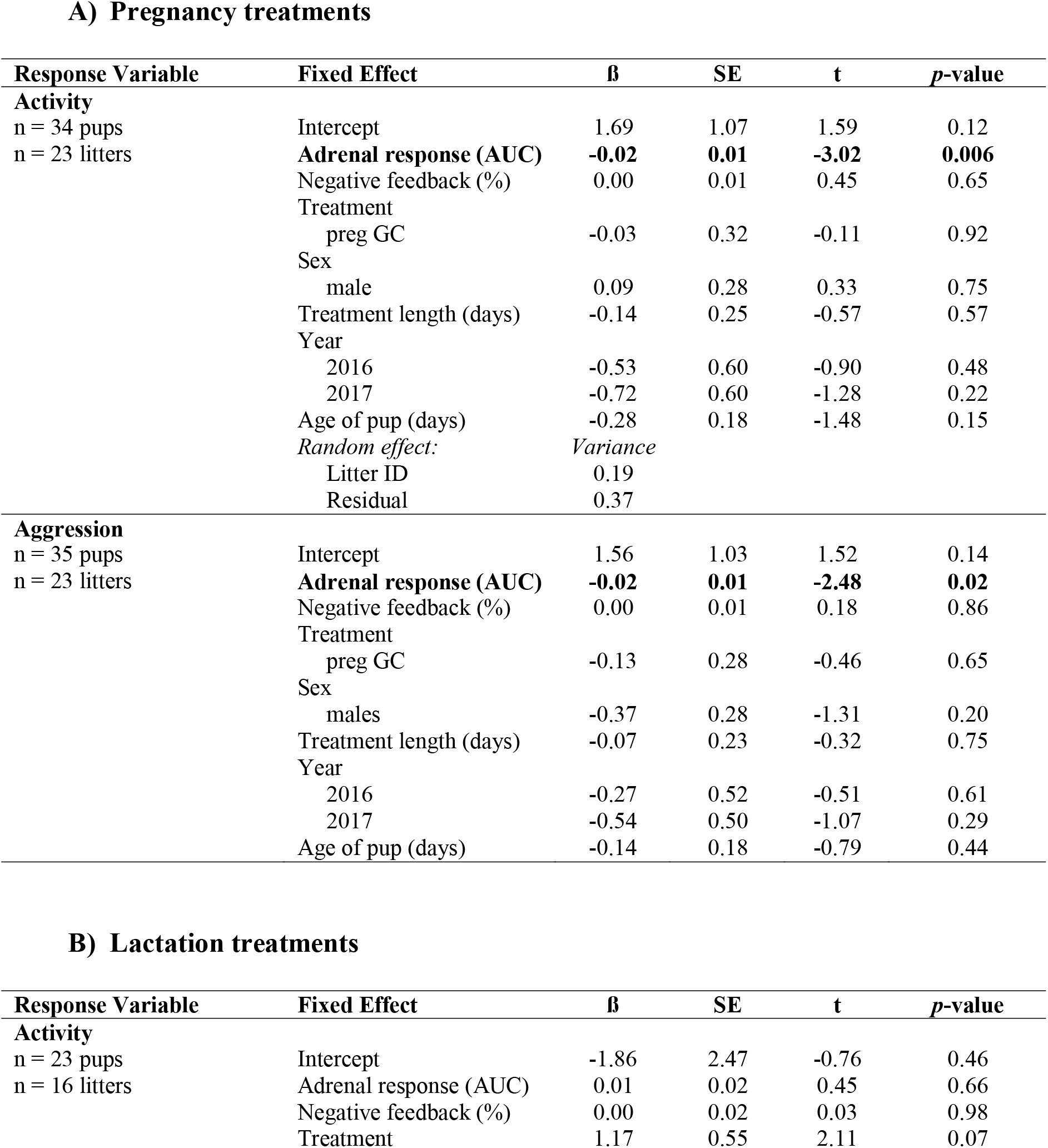
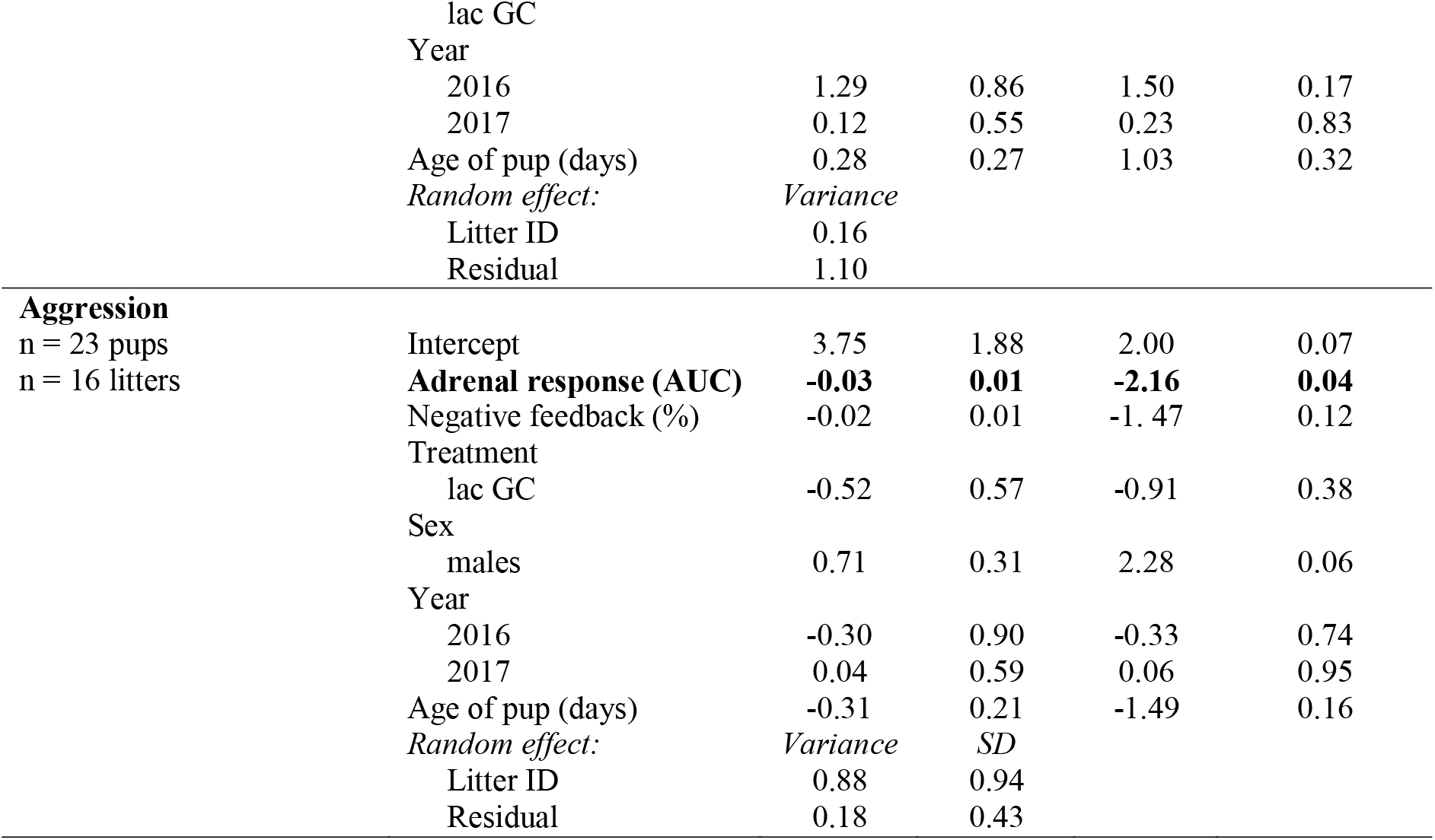
Full results of behavioral trait models. We ran four distinct models testing the effect of maternal GCs on activity and aggression in juvenile red squirrels. We ran separate models for the (a) pregnancy and (b) lactation treatment groups and the two behavioral traits. We standardized treatment length (days) across all data for the pregnancy models and age of pup (days) across all data. The comparison group for categorical variables is control treatment females from 2015. Bold font indicates statistical significance of *p* < 0.05.

**A) Pregnancy treatments**
| Response Variable | Fixed Effect | $\beta$ | SE | t | p-value |
| --- | --- | --- | --- | --- | --- |
| **Activity** |  |  |  |  |  |
| n = 34 pups | Intercept | 1.69 | 1.07 | 1.59 | 0.12 |
| n = 23 litters | **Adrenal response (AUC)** | **-0.02** | **0.01** | **-3.02** | **0.006** |
|  | Negative feedback (%) | 0.00 | 0.01 | 0.45 | 0.65 |
|  | Treatment |  |  |  |  |
|  | preg GC | -0.03 | 0.32 | -0.11 | 0.92 |
|  | Sex |  |  |  |  |
|  | male | 0.09 | 0.28 | 0.33 | 0.75 |
|  | Treatment length (days) | -0.14 | 0.25 | -0.57 | 0.57 |
|  | Year |  |  |  |  |
|  | 2016 | -0.53 | 0.60 | -0.90 | 0.48 |
|  | 2017 | -0.72 | 0.60 | -1.28 | 0.22 |
|  | Age of pup (days) | -0.28 | 0.18 | -1.48 | 0.15 |
|  | Random effect: | Variance |  |  |  |
|  | Litter ID | 0.19 |  |  |  |
|  | Residual | 0.37 |  |  |  |
| **Aggression** |  |  |  |  |  |
| n = 35 pups | Intercept | 1.56 | 1.03 | 1.52 | 0.14 |
| n = 23 litters | **Adrenal response (AUC)** | **-0.02** | **0.01** | **-2.48** | **0.02** |
|  | Negative feedback (%) | 0.00 | 0.01 | 0.18 | 0.86 |
|  | Treatment |  |  |  |  |
|  | preg GC | -0.13 | 0.28 | -0.46 | 0.65 |
|  | Sex |  |  |  |  |
|  | males | -0.37 | 0.28 | -1.31 | 0.20 |
|  | Treatment length (days) | -0.07 | 0.23 | -0.32 | 0.75 |
|  | Year |  |  |  |  |
|  | 2016 | -0.27 | 0.52 | -0.51 | 0.61 |
|  | 2017 | -0.54 | 0.50 | -1.07 | 0.29 |
|  | Age of pup (days) | -0.14 | 0.18 | -0.79 | 0.44 |

**B) Lactation treatments**
| Response Variable | Fixed Effect | $\beta$ | SE | t | p-value |
| --- | --- | --- | --- | --- | --- |
| **Activity** |  |  |  |  |  |
| n = 23 pups | Intercept | -1.86 | 2.47 | -0.76 | 0.46 |
| n = 16 litters | Adrenal response (AUC) | 0.01 | 0.02 | 0.45 | 0.66 |
|  | Negative feedback (%) | 0.00 | 0.02 | 0.03 | 0.98 |
|  | Treatment lac GC | 1.17 | 0.55 | 2.11 | 0.07 |
|  | Year |  |  |  |  |
|  | 2016 | 1.29 | 0.86 | 1.50 | 0.17 |
|  | 2017 | 0.12 | 0.55 | 0.23 | 0.83 |
|  | Age of pup (days) | 0.28 | 0.27 | 1.03 | 0.32 |
|  | *Random effect:* | *Variance* |  |  |  |
|  | Litter ID | 0.16 |  |  |  |
|  | Residual | 1.10 |  |  |  |
| **Aggression** |  |  |  |  |  |
| n = 23 pups | Intercept | 3.75 | 1.88 | 2.00 | 0.07 |
| n = 16 litters | **Adrenal response (AUC)** | **-0.03** | **0.01** | **-2.16** | **0.04** |
|  | Negative feedback (%) | -0.02 | 0.01 | -1.47 | 0.12 |
|  | Treatment |  |  |  |  |
|  | lac GC | -0.52 | 0.57 | -0.91 | 0.38 |
|  | Sex |  |  |  |  |
|  | males | 0.71 | 0.31 | 2.28 | 0.06 |
|  | Year |  |  |  |  |
|  | 2016 | -0.30 | 0.90 | -0.33 | 0.74 |
|  | 2017 | 0.04 | 0.59 | 0.06 | 0.95 |
|  | Age of pup (days) | -0.31 | 0.21 | -1.49 | 0.16 |
|  | *Random effect:* | *Variance* | *SD* |  |  |
|  | Litter ID | 0.88 | 0.94 |  |  |
|  | Residual | 0.18 | 0.43 |  |  |

**Figure 3.**
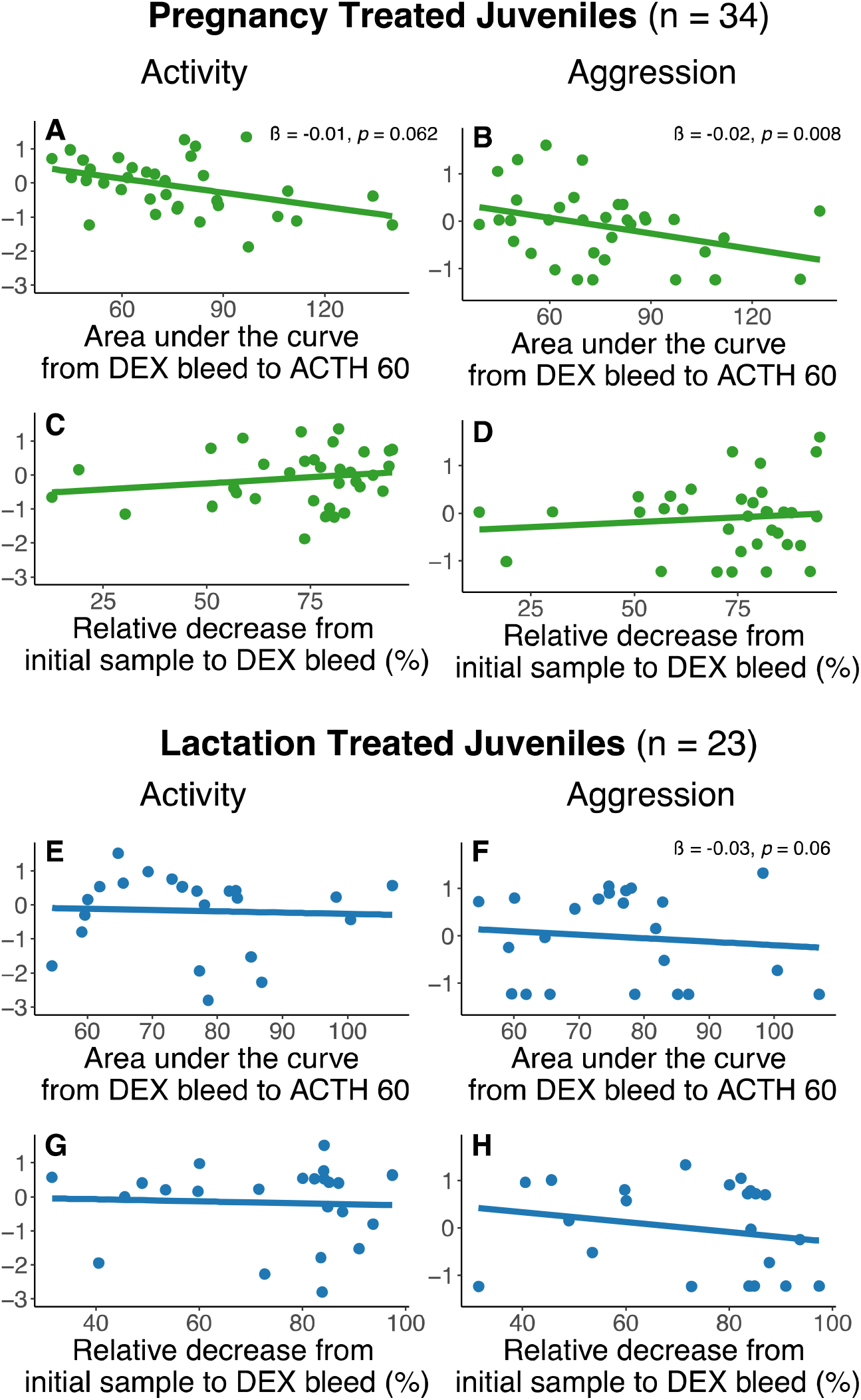
Relationships between behavioral traits and HPA axis dynamics. More active (A) and aggressive (B) juvenile red squirrels from the pregnancy treatment groups (both GC-treated and controls) exhibited smaller adrenal reactivity (area under the curve from the DEX bleed to the ACTH 60 min bleed; Table 4). More aggressive juveniles from the lactation treatment groups (both GC-treated and the controls) also exhibited smaller ACTH AUC (F; Table 4). Panels A-D include raw data from pregnancy treated juveniles (n = 34). Panels E-H include raw data from lactation treated juveniles (n = 23). Separate regression lines for each treatment group (GC-treated and controls) are not shown because there was no significant effect of treatment (Table 4). Regression coefficients and *p*-values are shown for models with *p*-values < 0.07. Panels A, C, E, and G show the relationship between activity and our two measures of HPA axis dynamics. Panels B, D, F, and H show the relationship between aggression and HPA axis dynamics.

**Figure 4.**
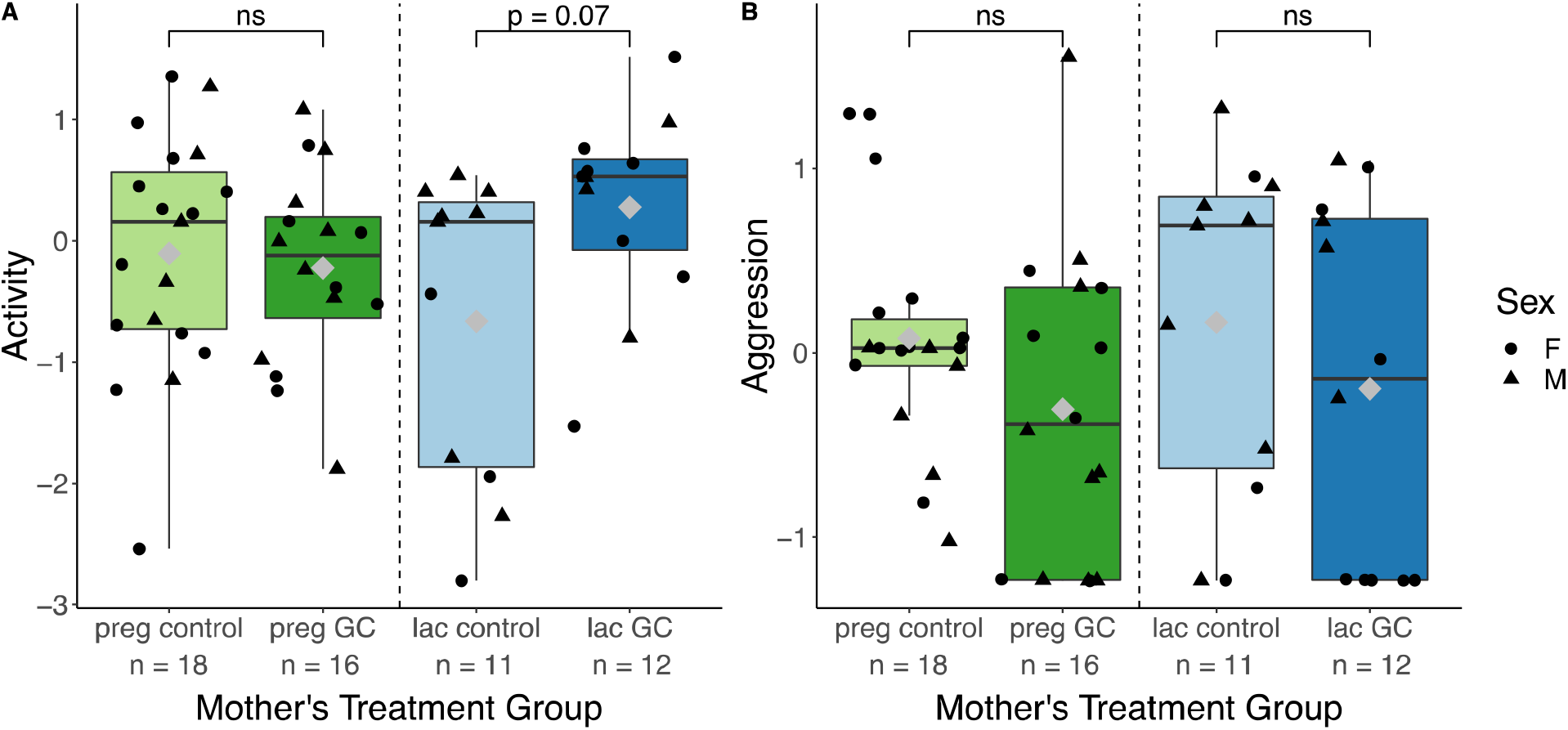
Effects of treatment group on activity and aggression. (A) Activity and (B) aggression were not significantly impacted by maternal treatment with GCs, though lac GC juveniles showed slightly higher levels of activity than controls (Table 4). Grey diamonds indicate the group means. Solid lines within the boxes indicate the group median. Data points represent individual juvenile red squirrels (preg control n = 18, preg GC n = 16, lac control n = 11, lac GC n = 12). The shape of the data point indicates the sex of the individual (circle = female red squirrel, triangle = male red squirrel). *P*-value of comparison between the corresponding control and GC treatment groups is indicated above the box plot if *p* < 0.08 (ns = not significant).

Among all juveniles, more aggressive individuals had a lower adrenal response to ACTH than less aggressive individuals (pregnancy treatment: ß = -0.02, *p* = 0.02; lactation treatment: ß = -0.03, *p* = 0.04; Table 4; Figure 3). In the lactation treatment group, males were slightly more aggressive than females, though this sex difference was not statistically significant (ß = 0.71, *p* = 0.06; Table 4) No other factors in the models predicted variation in aggression (Table 4).

### Relationship between behavioral traits and HPA axis activity

Treatment with exogenous GCs during pregnancy or lactation did not affect the relationship between activity and aggression (pregnancy treatment: ß = 0.06, *p* = 0.91; lactation treatment: ß = -0.34, *p* = 0.50; Table 5). Aggression was not predictive of activity in either pregnancy or lactation treated individuals, though the effect was in the expected direction, of more aggressive individuals exhibiting more active behavior based on previous studies with much larger sample sizes (pregnancy treatment: ß = 0.49, *p* = 0.21; lactation treatment: ß = 0.56, *p* = 0.15; Table 5; Figure 5). Treatment group, treatment length, juvenile age, juvenile sex, and year of experiment all showed no relationship with activity among pregnancy treated juveniles (Table 5). Contrary to the previous model for activity, in this model, which does not include HPA axis variables, lac GC juveniles showed higher activity levels than lac control juveniles (ß = 1.12, *p* = 0.046; Table 5).

**Table 5.** Full results of the relationship between behavioral traits models. We ran two distinct models testing the effect of maternal GCs on the relationship between activity and aggression among juvenile red squirrels. We ran separate models for the (a) pregnancy and (b) lactation treatment groups. We standardized treatment length across all data for the pregnancy model. The comparison group for categorical variables is control treatment females from 2015. Bold font indicates statistical significance of *p* < 0.05.

**A) Pregnancy treatments**
| Response Variable | Fixed Effect | $\beta$ | SE | t | p-value |
| --- | --- | --- | --- | --- | --- |
| Activity<br>n = 35 pups<br>n = 24 litters | Intercept | -0.05 | 0.44 | -0.13 | 0.90 |
|  | Aggression | 0.49 | 0.39 | 1.29 | 0.21 |
|  | Treatment |  |  |  |  |
|  | preg GC | -0.02 | 0.35 | -0.06 | 0.95 |
|  | Sex |  |  |  |  |
|  | male | 0.34 | 0.35 | 0.97 | 0.34 |
|  | Treatment length (days) | 0.04 | 0.28 | 0.14 | 0.89 |
|  | Year |  |  |  |  |
|  | 2016 | -0.08 | 0.52 | -0.16 | 0.88 |
|  | 2017 | -0.34 | 0.61 | -0.56 | 0.58 |
|  | Aggression*Treatment | 0.06 | 0.49 | -0.12 | 0.91 |
|  | Random effect: | Variance |  |  |  |
|  | Litter ID | 0.09 |  |  |  |
|  | Residual | 0.72 |  |  |  |

**B) Lactation treatments**
| Response Variable | Fixed Effect | $\beta$ | SE | t | p-value |
| --- | --- | --- | --- | --- | --- |
| Activity<br>n = 23 pups<br>n = 16 litters | Intercept | -0.97 | 0.57 | -1.70 | 0.11 |
|  | Aggression | 0.56 | 0.37 | 1.52 | 0.15 |
|  | Treatment |  |  |  |  |
|  | **lac GC** | **1.12** | **0.49** | **2.27** | **0.046** |
|  | Sex |  |  |  |  |
|  | male | 0.29 | 0.54 | 0.54 | 0.60 |
|  | Year |  |  |  |  |
|  | 2016 | 0.59 | 0.73 | 0.81 | 0.45 |
|  | 2017 | -0.09 | 0.51 | -0.17 | 0.87 |
|  | Aggression*Treatment | -0.34 | 0.49 | -0.70 | 0.50 |
|  | Random effect: | Variance |  |  |  |
|  | Litter ID | 0.04 |  |  |  |
|  | Residual | 1.05 |  |  |  |

**Figure 5.**
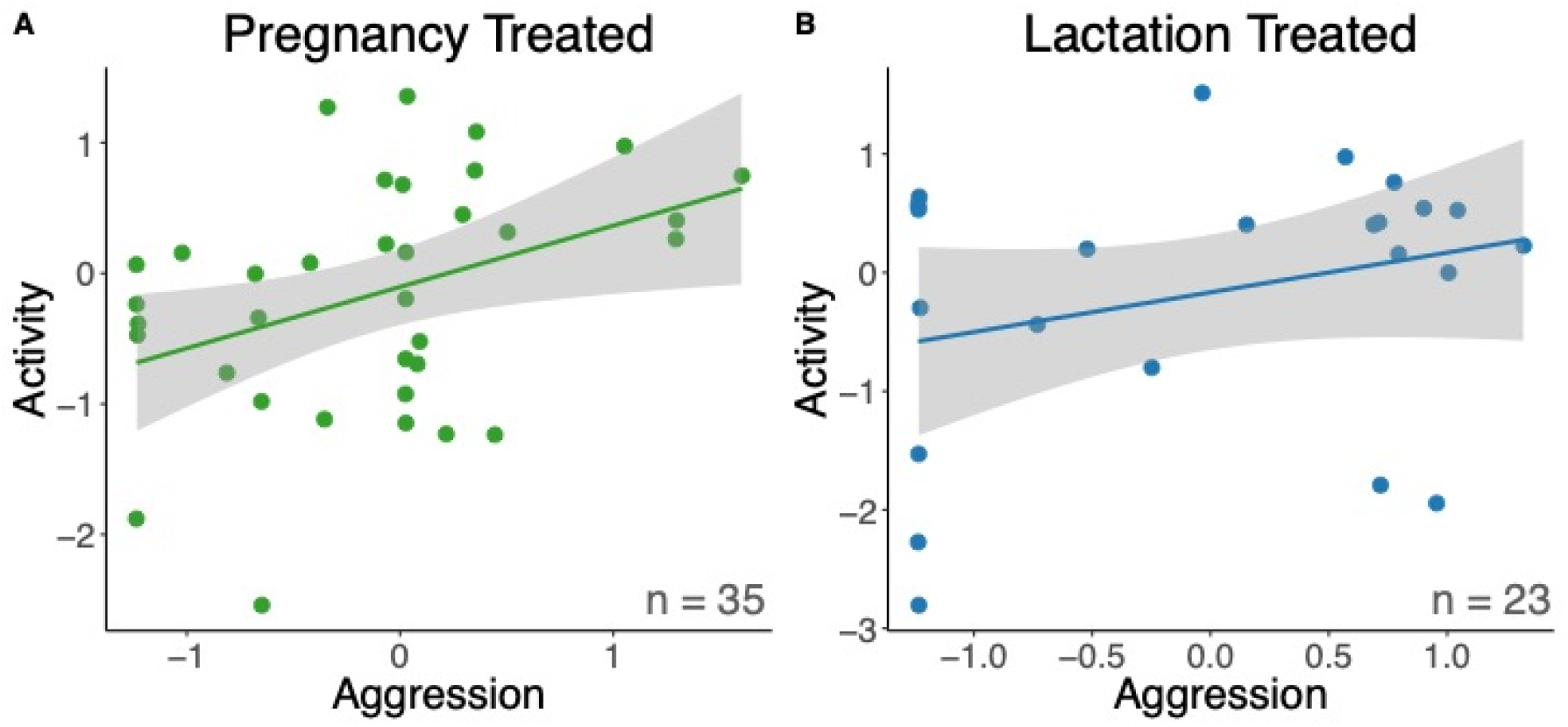
Relationship between activity and aggression. The relationship between the PCA scores for both behavioral traits (activity and aggression) was positive, though not statistically significant (Table 5), for the juvenile red squirrels from both pregnancy (A) and lactation (B) treated mothers. There was no significant interaction between treatment and aggression on activity levels (Table 5), and so GC and control individuals for each period (pregnancy or lactation) were combined and only the main effect of aggression was plotted. Grey shaded areas indicate 95% confidence intervals of the fitted regression line. Green data points represent individual juvenile squirrels from pregnancy treated mothers (n = 35) and blue data points represent individual juvenile squirrels from lactation treated mothers (n = 23).

## Discussion

Overall, our experimental increase in maternal GCs during pregnancy or lactation had only subtle effects on behavioral traits of juvenile offspring. Activity was slightly higher in the lac GC treatment group compared to controls, but there were no statistically significant differences between the pregnancy treatment groups, and aggression was not affected by the GC treatments. Exogenous GCs provided to pregnant or lactating females did not affect their offspring’s HPA axis response to dexamethasone (negative feedback) nor ACTH (adrenal reactivity). However, we did find a relationship between one measure of HPA axis activity and both behavioral traits of the juvenile squirrels we tested. Less active squirrels from the pregnancy treatment groups were more responsive to ACTH, but we did not find this relationship among squirrels in the lactation treatment groups. Similarly, less aggressive squirrels were more responsive to ACTH among all juveniles from all of the pregnancy and lactation treatment groups. The negative feedback response of the HPA axis was unrelated to activity and aggression.

Contrary to our findings, a recent meta-analysis including 39 studies across 14 vertebrate species found an overall positive relationship between prenatal stress and offspring GC levels, with a particularly strong effect for experimental studies compared to observational ones (Thayer et al., 2018). This meta-analysis also found a stronger effect of prenatal stress on the negative feedback of the HPA axis than baseline or peak GC response to a stressor (Thayer et al. 2018). It is important to note this meta-analysis includes studies with either administration of exogenous GCs or general prenatal stress exposure, so it does not directly address the impact of exogenous GCs in particular (Thayer et al. 2018). There are multiple non-mutually exclusive potential explanations for why our experiment did not follow these trends. Two potential explanations involve potential bias in our experimental design. First, inevitably not all offspring from the pre- and post-natal GC supplementation experiment survived to weaning (van Kesteren et al., 2019), and potentially not all offspring from the experiment stayed within our study range (though this is unlikely given relatively short natal dispersal distances, see: Berteaux and Boutin, 2000; Cooper et al., 2017; Kerr et al., 2007; Martinig et al., 2020), both of which could have resulted in a survivor bias in our results. Second, we may have had selection biases in our trapping success rate. Despite extensive trapping efforts, differences in behavior among offspring may have resulted in a reduction in our ability to trap less aggressive, less active squirrels, and led to underestimation of effect sizes, which would reduce our power to detect any correlations and treatment group effects (Carter et al., 2012; Kelley et al., 2015). However, based on the number of juveniles alive at weaning (determined via yearly population census, trapping, and behavioral observations), we actually included a majority of the juveniles born to GC/control treated squirrels (68%) in this study. Based on census, trapping, and behavioral observations, 32% of juveniles born to a treated squirrel were alive at the time of sampling but were never trapped for sampling. Some of these juveniles were purposely excluded from sampling to avoid collecting data for more than two individuals per litter.

One proximate mechanism potentially limiting our detection of effects of maternal GCs in our experiment could be the ongoing neural pruning of important brain structures responsible for consistent behavioral traits and reactions to external stressors between early life and adolescence (Groothuis and Trillmich, 2011; Spear, 2000). In other words, the effects of the GC manipulation may not have lasted into adolescence due to this neural pruning, or our manipulation was not sufficiently long enough to cover this critical developmental window. The HPA axis of weaned juveniles shows a functional response to both ACTH and DEX, but may still be fine-tuned as the squirrel matures. In this environment, it may be adaptive to adjust behavioral phenotypes throughout development (Kelley et al., 2015), and by testing these offspring at the stage of weaning, their behavior and HPA axis may have been shaped more by their own phenotypes and personal experiences than their early-life environment (Nettle and Bateson, 2015).

In this region, predation risk, resource availability, and conspecific competition fluctuate between an individual’s birth and first breeding season (Dantzer et al., 2013; Hendrix et al., 2020; McAdam and Boutin, 2003; Taylor et al., 2014). However, environmental factors known to be influential in the life history of red squirrel in this region, specifically resource availability and conspecific density, fluctuate on a yearly basis due to the annual reproductive cycle of red squirrels and white spruce trees (Dantzer et al., 2013; McAdam and Boutin, 2003; Taylor et al., 2014). Therefore, we would predict most aspects of the parental environment would be consistent with the environment the offspring will experience as they wean and establish their territory before overwintering, which is the time point at which we tested them. However, it is possible that too much sensitivity to the early life environment may be maladaptive for a species in a highly variable environment. Theoretically, the short developmental of red squirrels window could be affected by short-term stochastic processes that we are currently unaware of that result in a mismatch between the parental environment and offspring environment and, in that case, it would be maladaptive for offspring to be attuned to cues subject to short-term stochastic processes (Langenhof and Komdeur, 2018). In essence, the world the developing juveniles experienced may have contradicted the environmental information that was conveyed by the cues of maternal GCs, and therefore the maternal cues may have been “overwritten” by the immediate cues (Leimar and McNamara, 2015).

In addition to testing maternal GC treatment effects, our results also provide evidence supporting the original unidimensional coping styles model (Koolhaas et al., 1999) which posits more active, aggressive individuals should exhibit a decreased physiological stress response, specifically decreased HPA axis activation. Previously, we tested the unidimensional (Koolhaas et al., 1999) and two-tier coping style (Koolhaas et al., 2010) models in adult red squirrels and found no relationship between the physiological and behavioral stress responses, thus supporting the two-tier model (Westrick et al., 2019). While the conclusions of our two studies are different, we believe the studies are not directly comparable. There are two important ways in which our two studies differ.

First, in our previous study, although we used the same behavioral trial methods, we used fecal glucocorticoid metabolite concentrations as an integrative measurement of HPA axis activity (Westrick et al., 2019). It is possible that the relationship between HPA axis activity and behavioral traits was not detectable on that broad of a scale, whereas measuring plasma cortisol in response to a standardized challenge could provide more relevant measurement of the acute HPA axis response. However, research on snowshoe hares (*Lepus americanus*) suggests that higher fecal glucocorticoid metabolite concentrations correspond to a smaller decrease in plasma cortisol concentrations in response to DEX and a larger response ACTH (Sheriff et al., 2010).

Second, our previous study looked at the behavioral and physiological stress response in adult (>1 year old) squirrels, whereas our current study involves juvenile red squirrels around the period of weaning (∼70 days old). Previous work has shown that these behavioral traits “regress to the mean” in this study system, meaning there is less variation and a lack of individuals at the extremes of activity and aggression in the adult population compared to the juvenile population (Kelley et al., 2015). This could mean the wide range in behavioral stress responses in the juvenile population of red squirrels is more consistent with the previous laboratory studies of other species and selection lines for individuals at the extreme ends of the proactive-reactive spectrum (which often find support for the unidimensional model: Koolhaas et al. 1999; Westrick et al. 2019) compared to the adult population of red squirrels. It is also possible that only individual juveniles with low covariance between the behavioral and physiological stress responses were able to adaptively respond to conditions, due to the two stress responses being decoupled. Therefore, the surviving adult population is less likely to exhibit high covariance between HPA axis activity and behavioral traits. Additionally, young animals need to be adapted to the experience of that life stage to survive to adulthood, which may require a different set of adaptations (Groothuis and Trillmich, 2011). We believe these two findings in the same study system serve as further evidence of a need for a more generalizable model of the relationship between the behavioral and physiological stress response, as highlighted in Westrick et al. (2019).

Our study is one of the first to use an experimental manipulation to understand the effects of the maternal environment, as encoded through GCs, on the ontogeny of personality traits in a wild population. Given the context of our extensive knowledge around the selection pressures acting on these specific behavioral traits (Boon et al., 2007, 2008; Cooper et al., 2017; Kelley et al., 2015; Taylor et al., 2012, 2014) and our knowledge of how maternal GCs shape specific aspects of early development in red squirrels (Dantzer et al., 2013, 2020), this study system provided the ideal opportunity to test the developmental role of maternal GCs in personality in a natural environment (Groothuis and Trillmich, 2011; Langenhof and Komdeur, 2018). The fact that we did not find a substantial effect of pre- or post-natal GCs on HPA axis activity and behavior of juvenile squirrels may indicate maternal GCs are not an ecologically relevant cue for the development of these traits. It may be that, as a short-lived animal in a highly-variable environment, juvenile squirrels are less sensitive to maternal GC cues despite the proposed links between maternal GCs, the HPA axis development, and behavior. Based on our results, maternal GCs do not adaptively drive developmental plasticity of behavior or the phenotypic correlations between behavior and HPA axis dynamics in recently weaned red squirrel juveniles.

## Acknowledgements

We thank Agnes MacDonald, her family, and Champagne and Aishihik First Nations for allowing us to conduct our work within their traditional territory. We thank Monica Cooper, Zach Fogel, Claire Hoffmann, Noah Israel, Sean Konkolics, Laura Porter, Matt Sehrsweeny, Sam Sonnega, Jess Steketee, and Dylan Yaffy for their assistance in field data collection. We thank Brie Coleman, Deirdre McGovern, Austin Rife, and Meg Ryan for their assistance in scoring videos. This is publication XX of the Kluane Red Squirrel Project.

## Competing interests

No competing interests declared.

## Funding

This work was supported by American Society of Mammalogists to SEW; University of Michigan to SEW and BD; National Science Foundation (IOS-1749627 to BD); and Natural Sciences and Engineering Research Council to SB, AGM, and JEL.

## Data availability

All data are available from figshare: temporary private link to be updated with public DOI (https://figshare.com/s/9f5846adda84f6e7bb02)

## Ethical statement

All work was conducted under animal ethics approvals from University of Michigan (PRO00005866).

